# Licensing Microgels Prolong the Immunomodulatory Phenotype of Mesenchymal Stromal Cells

**DOI:** 10.1101/2022.07.05.498567

**Authors:** Matthew Patrick, Ramkumar T. Annamalai

**Affiliations:** Department of Biomedical Engineering, University of Kentucky, Lexington, KY 40506

**Keywords:** Microgel, Interferon Gamma, Bone, Mesenchymal Stem cells, Licensing

## Abstract

Mesenchymal stromal cells (MSC) are sensors of inflammation, and they exert immunomodulatory properties through the secretion of cytokines and exosomes and direct cell-cell interactions. MSC are routinely used in clinical trials and effectively resolve inflammatory conditions. Nevertheless, inconsistent clinical outcomes necessitate the need for more robust therapeutic phenotypes. The immunomodulatory properties of MSC can be enhanced and protracted by priming (aka licensing) them with IFNγ and TNFα. Yet these enhanced properties rapidly diminish, and prolonged stimulation could tolerize their response. Hence a balanced approach is needed to enhance the therapeutic potential of the MSC for consistent clinical performance. Here, we investigated the concentration-dependent effects of IFNγ and TNFα and developed gelatin-based microgels to sustain a licensed MSC phenotype. We show that IFNγ treatment is more beneficial than TNFα in promoting an immunomodulatory MSC phenotype. We also show that the microgels possess integrin-binding sites to support MSC attachment and a net positive charge to sequester the licensing cytokines electrostatically. Microgels are enzymatically degradable, and the rate is dependent on the enzyme concentration and matrix density. Our studies show that one milligram of microgels by dry mass can sequester up to 641 ± 81 ng of IFNγ. Upon enzymatic degradation, microgels exhibited a sustained release of IFNγ that linearly correlated with their degradation rate. The MSC cultured on the IFNγ sequestered microgels displayed efficient licensing potential comparable to or exceeding the effects of bolus IFNγ treatment. When cultured with proinflammatory M1-like macrophages, the MSC-seeded on licensing microgel showed an enhanced immunomodulatory potential compared to untreated MSC and MSC treated with bolus IFNγ treatment. Specifically, the MSC seeded on licensing microgels significantly upregulated *Arg1, Mrc1,* and *Igf1*, and downregulated *Tnfa* in M1-like macrophages compared to other treatment conditions. These licensing microgels are a potent immunomodulatory approach that shows substantial promise in elevating the efficacy of current MSC therapies and may find utility in treating chronic inflammatory conditions.

## 1. Introduction

Mesenchymal stromal cell (MSC) therapy has attracted clinicians and researchers as a viable treatment option for many inflammatory and immune diseases. MSC are a potent immunomodulatory tool that can regulate innate and adaptive immune responses through direct cell-cell and paracrine signaling^1^. Their potential to modulate disease states has been widely recognized, and there were over 900 clinical trials between 2004 and 2018 in which MSC were the primary disease-modifying agent^2^. Further, at least 10 MSC-based therapies are approved globally for inflammatory and immune diseases such as graft-versus-host disease, Crohn’s disease, amyotrophic lateral sclerosis (ALS), and myocardial infarction^3^. Although their effectiveness in treating certain diseases has been well documented, the outcomes are inconsistent and vary with the treatment timing and the inflammatory milieu^4^.

MSC are sensors and modifiers of the body’s immune response, regulating the expression of specific cytokines, chemokines, and growth factors. During low inflammatory states, MSC adopt a quiescent phenotype with low prostaglandin E2 (PGE2) and HLA-G expression that are vital for sustaining T_H_1 and T_H_17 proliferation^5, 6^. But during inflammation, MSC are activated or adopt a ‘licensed’ phenotype resulting in increased expression of immunomodulatory factors, including indolamine-2,3-dioxygenase (IDO, in humans), nitric oxide (NO, in mice), PGE2, TNFα-stimulated gene-6 (TSG6), transforming growth factor-β (TGF-β), IL-10, and IL-1RA^7^. These factors suppress T_H_1 and T_H_17 cells and promote immunosuppressive T_regs_, thereby modulating the local inflammatory macrophages toward a pro-healing phenotype^8^. It should be noted that the degree of MSC activation and subsequent modulation is context-specific and varies with the changing disease state. Although these disease-modifying properties are beneficial for treating benign inflammatory conditions, their usefulness in chronic conditions is restrictive and inconsistent.

The extensive tissue damage and weakened healing response seen in many inflammatory pathologies result from an abnormally heightened or prolonged inflammatory state. In the clinic, MSC are administered locally or systemically which are then licensed by the disease-invoked inflammatory milieu to provide an immunomodulatory response. When administered locally, they are rapidly cleared from the site^3, 9^. Hence, their therapeutic response primarily happens during circulation or while entrapped in a capillary bed of the lungs before being cleared from the body^10^. In conditions characterized by chronic inflammation, such as diabetes and arthritis, the therapeutic effects may not persist long enough for proper treatment. Further, the presence of an inflammatory milieu is not guaranteed, and even when pre-licensed, their therapeutic phenotype rapidly diminishes. These factors contribute to the overall inconsistent outcomes of MSC therapies. Addressing these limitations can help improve the efficacy and the clinical relevance of MSC-based therapies.

Strategies for pre-licensing MSC with inflammatory cytokines are being studied to promote a consistent and sustained therapeutic phenotype. Pre-licensing with either IFNγ or TNFα has also been shown to improve the intensity of their immunomodulatory effects. Stimulation of the MSC IFNγ receptor or TNFα receptor activates the JAK/STAT pathway or NFκB pathway respectively and leads to the subsequent regulation of the immunomodulatory factors^11^. In animal models, IFNγ-licensed MSC are able to ameliorate inflammatory diseases like GvHD and colitis^12, 13^. Similarly, TNFα-licensed MSC exhibit therapeutic effects in a mouse model of inflammatory bowel disease^14^. Hence, localized and sustained release of these licensing factors could prolong and augment the immunomodulatory effects of MSC.

Material-based systems are used widely in applications that require a sustained release of biochemical factors and deliver therapeutic cells. For instance, bone tissue engineering approaches that release osteoinductive factors have already seen clinical application^15^. Material-based systems are customizable and suitable to augment the therapeutic efficacy of cells and biomolecules. Here, we used a modular microgel system that can sequester licensing cytokines while providing a suitable surface for MSC attachment and proliferation. The enzymatic degradation of the microgel matrix by the MSC releases the licensing factors locally to sustain an immunomodulatory phenotype of MSC. The microgels are composed of biocompatible gelatin polypeptides containing RGD motifs facilitating integrin-mediated cell attachment^16^. Further, the polypeptides are crosslinked with genipin, a naturally occurring crosslinker with good cytocompatibility, to enhance their mechanical strength^17^. This crosslinking depletes the positively charged amines of the lysine residues imparting a net negative charge on the crosslinked polypeptides^18^, allowing ionic sequestration of biomolecules with a positively charged domains^19^. MSC can be cultured on the surface of these microgels sequestered with cytokines on the core matrix and injected into the injury site in a non-invasive manner. Attachment to the microgel will increase MSC retention at the injection site, while the sequestered cytokines will provide local and sustained delivery to maintain a licensed phenotype. Additionally, the microgel’s unique physical characteristics, including the shape and stiffness, make them ideal for promoting osteogenesis of MSC and subsequent integration with the host tissue.

Here, we characterized the physical and biochemical properties of the microgels and demonstrated their ability to ameliorate inflammatory conditions. We show that the microgels efficiently sequester IFNγ and release it upon degradation while readily supporting MSC attachment. We also show that the localized release of IFNγ promotes an immunomodulatory MSC phenotype efficiently. Further, we confirm their immunomodulatory phenotype through macrophage coculture studies. In addition, we evaluated the osteogenic potential of the microgels to show that they naturally direct MSC toward an osteogenic line over a prolonged culture period. Overall, our microgels show substantial promise in elevating the efficacy of current MSC therapies and demonstrate an improved option to address persistent chronic inflammatory diseases.

## 2. Methods

### 2.1. Microgel fabrication

Microgels were fabricated as previously described^19^. Briefly, gelatin derived from porcine skin (type A, 300 bloom, Sigma) was dissolved in deionized (DI) water to make a 6 wt% stock solution. The stock solution was dispensed dropwise into a polydimethylsiloxane (PDMS, viscosity = 100 cS, Clearco Products Co., Inc.) bath heated to 37°C and stirred by a propeller impeller at 500 rpm. After 5 min of emulsification, the PDMS bath was cooled with an ice bath for 30 min. They were then collected through a series of 275 g centrifugation and wash steps using Dulbecco phosphate buffer solution (DPBS, Invitrogen) supplemented with 1% TWEEN 20 (Sigma). The microgels were then crosslinked by adding 1 wt% genipin (Wako Chemicals) and left in an auto rotator for 48 hours. The microgels were then washed with 100% ethanol to stop the crosslinking and remove excess genipin and swelled in DPBS for 24 hours. Finally, they were sonicated to remove aggregates and filtered to the desired size range using nylon mesh sieves.

### 2.2. Size distribution and swelling rate

Bright-field imaging of microgels was done to measure the diameter and determine the size distribution of microgels. Microgels were dehydrated using 100% ethanol, flash-frozen in liquid nitrogen, and lyophilized overnight. Dehydrated microgels were rehydrated in DPBS, monitored using an inverted microscope, and recorded periodically. The images were then processed and analyzed using Fiji (ImageJ) software (National Institutes of Health) to determine the size and swelling rates.

### 2.3. Enzymatic degradation

A stock solution was prepared by adding a precise amount of dehydrated and lyophilized microgels in 1X DPBS (stock = 0.67 mg/mL). 200 μL of the sock solution (~130 μg/well) was dispensed in a 96 well plate, and an equal volume of collagenase solution (Worthington Biochemical) was added to the wells at different concentrations and sealed with a clear plate cover to prevent evaporation. The fluorescence of the supernatant was then measured at Ex/Em 590/620 every 5 min for 48 hours to assess microgel degradation rates.

### 2.4. Loading and release characteristics of microgels

1 μg of IFNγ suspended in 5 μL of DPBS was added to 1 mg of lyophilized microgels and placed in an incubator for 24 hours. After loading, they were washed with 0.1 wt% albumin in DPBS to remove unbound IFNγ. To study the release kinetics, the loaded microgels were suspended in 1 mL of 100 U/mL collagenase solution and kept at 37°C until completely degraded. The supernatant was collected periodically and analyzed using an IFNγ ELISA kit to measure the release of IFNγ from the degrading microgel. The corresponding degradation rates of the microgels were also measured fluorometrically at Ex/Em 590/620.

### 2.5. Isolation of primary adipose-derived mesenchymal stem cells

Inguinal fats pads were isolated from adult mice (wildtype C57BL/6J – JAX stock #000664 and ACTb-EGFP expressing C57BL/6-Tg(CAG-EGFP)1Osb/J, JAX stock #003291), washed in serial dilutions of betadine and sterile 1X DPBS. The washed tissues were then finely minced with sterile scissors, and suspended in 0.1 wt% collagenase solution, and incubated at 37°C with constant agitation for 30 min or until most of the tissue was degraded. An equal amount of Dulbecco’s modified eagle’s medium (DMEM, Gibco) with 10% fetal bovine serum (FBS, Gibco) was then added to neutralize the collagenase and filtered through a 100 μm cell strainer. The single-cell suspension was then centrifuged at 250 g for 5 min, supernatant aspirated, and the cell pellet was resuspended in DMEM+10% FBS and cultured in a 6 well plate. After 24 hours, the wells were washed twice with warm 1x DPBS to remove non-adherent cells and debris leaving the adherent mesenchymal stromal cells (MSC). The cells were culture-expanded until the 5^th^ passage and used for experiments.

### 2.6. MSC culture and licensing

Primary MSC were cultured in MSC growth media containing DMEM+10% FBS+ antibiotics and antimycotics (Gibco). An IC-21 murine macrophage cell line (ATCC) was acquired, cultured in expansion media (RPMI (Gibco) + 10% FBS + antibiotics and antimycotic), and used for coculture studies. All cells were maintained at 37°C in a CO2 incubator. To license MSC, they were cultured until ~80 confluent in growth media and replaced with licensing media, i.e., growth media supplemented with the cytokines (TNFα, IFNγ, or both), for 24 hours. For coculture studies, macrophages were cultured in corresponding growth media until ~80% confluent and then polarized for 24 hours using cytokines (50 ng/mL LPS and 25 ng/mL IFNγ) suspended in growth media.

### 2.7. Quantitative gene expression

For gene expression analysis, the samples were collected in TRIzol™ reagent (Invitrogen), and the whole RNA was isolated according to the manufacture’s protocol. A SuperScript™ III Platinum™ One-Step qRT-PCR Kit (ThermoFisher) and TaqMan probes (ThermoFisher) were used to analyze gene expression profiles using a QuantStudio 3 real-time PCR system (ThermoFisher). The following TaqMan probes were used for our studies: Gapdh (Mm99999915_g1) as a housekeeping control, Alp (Mm00475834_m1), Bglap (Mm03413826_mH), Col1a1 (Mm00801666_g1), Ibsp (Mm00492555_m1), Sp7 (Mm04209856_m1), Spp1 (Mm00436767_m1), Runx2 (Mm00501584_m1) for MSC osteogenic analysis. Il-1β (Mm00434228_m1), Nos2 (Mm00440502_m1), Ccl2 (Mm00441242_m1), Tnfα (Mm00443260_g1), Lgals9 (Mm00495295_m1), Tgfβ (Mm01178820_m1), Il-6 (Mm00446190_m1), Ptgs2 (Mm00478374_m1), Pparγ (Mm01184322_m1), Arg1 (Mm00475988_m1), Cxcl2 (Mm00436450_m1), Mrc-1 (Mm00485148_m1), Il-12β (Mm00434174_m1), Igf1 (Mm00439560_m1) and, Chil3 (Mm00657889_mH). Gene expression data were presented as heatmaps showing foldchange (different treatment) or Z-scores (concentration and time-dependent change).

### 2.8. Cell proliferation

Cell proliferation was analyzed by quantifying the total DNA content of samples using Quant-iT™ PicoGreen™ dsDNA Assay Kit (ThermoFisher). Lambda DNA standard was used for generating a calibration curve. Briefly, samples were collected at the specified time points and homogenized through sonication in 1X Tris-HCl buffer and centrifuged at 10,000 g for 10 min. The supernatant was then mixed with PicoGreen, incubated in the dark for 5 min, and the fluorescence was read at Ex/Em wavelength of 485/528 nm.

### 2.9. Orthocresolphthalein complexone (OCPC) assay for calcium deposition

The o-cresolphthalein complex one (OCPC) method was used to quantify calcium deposition by the MSC on microgels as previously described^20^. Briefly, the samples were homogenized through sonication in 1X Tris-HCl buffer and centrifuged at 10,000 g for 10 min. The supernatant was then collected and mixed with a working solution containing 5% ethanolamine/boric acid buffer (Sigma), 5% o-cresolphthalein (Sigma), and 2% hydroxyquinoline (Sigma) in DI water. The mixture was incubated at room temperature in the dark for 10 min, and the absorbance was read at 575 nm. CaCl2 (Sigma) dissolved in DI water at different concentrations served as standards.

### 2.10. Sircol™ soluble collagen assay

The collagen deposition by the MSC was quantified using a Sircol soluble collagen assay kit (Biocolor). Briefly, the samples were homogenized through sonication in 1X Tris-HCl buffer and centrifuged at 10,000 g for 10 min. The supernatant was collected, and 20 μL of each sample was brought up to 100 μL with Tris-HCL buffer and then mixed with 0.5 mL of Sircol dye reagent. Samples were mixed for 30 min, centrifuged at 12,000 RPM for 10 min, the supernatant was decanted, and 750 μL of acid-salt wash reagent was added to the pellet. The suspension was mixed and centrifuged again at 12,000 RPM for 10 min, the supernatant was decanted, and 250 μL of alkali reagent was added to the pellet. This suspension was vortexed and incubated for 5 min at room temperature. The absorbance of the final solution was read at 555 nm. Rat-tail collagen was used to create a standard curve.

### 2.11. Alkaline phosphatase activity assay

A commercially available ALP activity fluorometric assay kit (BioVision) was used according to the manufacturer’s protocol. Briefly, the samples were homogenized through sonication in Tris-HCl buffer and centrifuged at 10,000 g for 10 min. Then 100 μL of the supernatant was mixed with 20 μL of 0.5 mM 4-methylumbelliferyl phosphate (MUP) substrate and incubated at 25°C in the dark for 30 min. Finally, 20 μL of stop solution was added to the mixture, and fluorescence was measured at Ex/Em of 360/440 nm. Samples of 0.1-0.5 nmol of MUP substrate with purified ALP enzyme were used to create a standard curve.

### 2.12. Immunofluorescence staining and confocal imaging

Samples were fixed in a 10% aqueous buffered zinc formalin (Z-Fix, Anatech) overnight, washed with 1X DPBS, and briefly incubated in a 0.1% Triton X-100 (Sigma) in 1X DPBS solution for 5 min before staining. Phalloidin conjugate (AF488, ThermoFisher) was used to stain the F-actin of the cell cytoskeleton and DAPI (Invitrogen) for staining the nucleus. Samples were stained for 30 mins in the dark and washed with 1X DPBS. Z-stacks of the stained samples were acquired using a Nikon A1R confocal microscope and processed using Fiji (ImageJ) software (NIH).

### 2.13. Electron microscopy

Samples were fixed with a 3% glutaraldehyde in sodium cacodylate buffer overnight, washed with DI water, and incubated in 2% Osmium Tetroxide solution for 16 hours. Samples were washed with DI water and dehydrated using ethanol series. Samples were then flash-frozen in liquid nitrogen and lyophilized overnight. The dried samples were sputter-coated with a 5 nm thick layer of platinum using Leica Ace 500 sputter coater and imaged using an FEI Quanta 250 environmental scanning electron microscope (SEM).

### 2.14. Statistics

All measurements were performed at least in triplicate. Data are plotted as means with error bars representing the standard deviation. The Pearson correlation coefficient (r) was used to evaluate the linear correlation between two variables. One and Two Way ANOVA were used to identify significance within groups, and Holms-Sidak pairwise comparisons followed to identify which groups were significant.

## 3. Results

### 3.1. Fabrication and characterization of microgels

Microgels were fabricated, crosslinked for 48 hours in genipin, and separated to the desired size range of ~150 μm. This size range was chosen to provide sufficient surface for MSC attachment and spreading, while maintaining injectability for minimally invasive applications. The spherical morphology and homogeneous size distribution of the microgels (**Fig.1 A-C**) allow them to conform to fill irregular wounds and articulating spaces. When suspended in collagenase solution, the microgels exhibited a concentration-dependent degradation (**Fig. 1D**). Similarly, when they were cultured with macrophages, a time-dependent degradation was seen (**Fig. 1E**). No degradation was seen in acellular control groups. The crosslinked microgel matrix possesses a net negative charge, as shown in the previous work^19^, enabling them to sequester biomolecules with a positive charge. Since IFNγ has an isoelectric point *(pI)* of ~8.7^21^, they are suitable for ionic-complexation and sequestration in our microgels. We achieved a maximum loading of 641 ± 81 ng of IFNγ per mg of lyophilized microgel. The release of IFNγ from the microgels depended on enzymatic degradation, and the release rate linearly correlated with the degradation rate (**Fig. 1F**).

**Figure 1.**
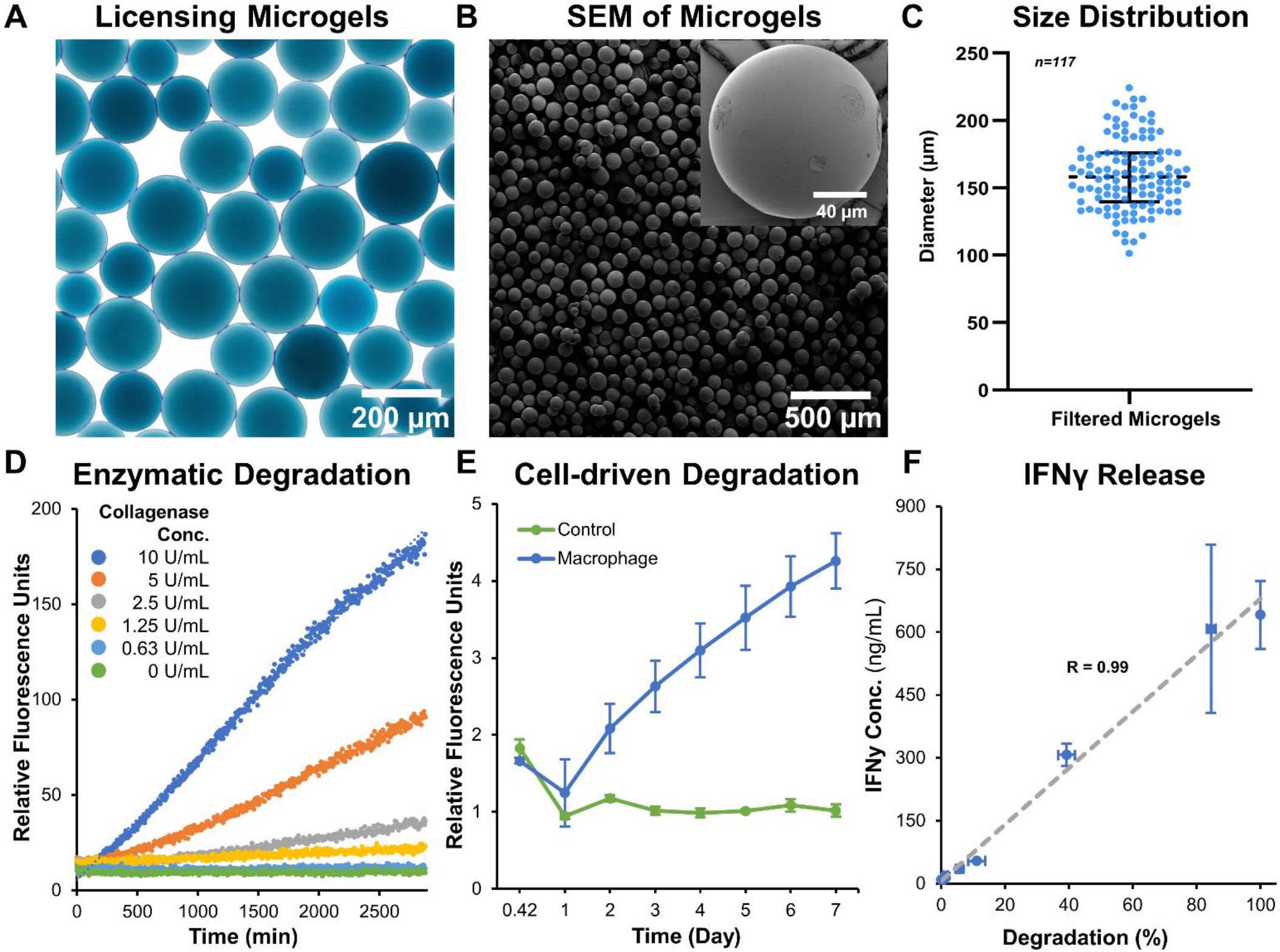
Microgel fabrication and characterization. **A)** Bright-field image of sorted microgels show homogenous size distribution. **B, C)** Scanning electron micrographs of dehydrated microgels show smooth surface and spherical morphology. **D)** Microgels exhibit a concentration-dependent degradation when incubated with collagenase solution. **E)** A similar time-dependent degradation was seen when cultured with macrophages. **F)** The release rate of IFNγ linearly correlated with the enzymatic-degradation of microgels (n =4).

The microgels also exhibit rapid swelling when suspended in an aqueous medium (**Fig. 2A**). The rate of hydration was rapid initially and plateaued within minutes to achieve an equilibrium hydration condition (**Fig. 2B**). An average percent increase in volume swelling ratio of ~400% was achieved in ~5 minutes.

**Figure 2.**
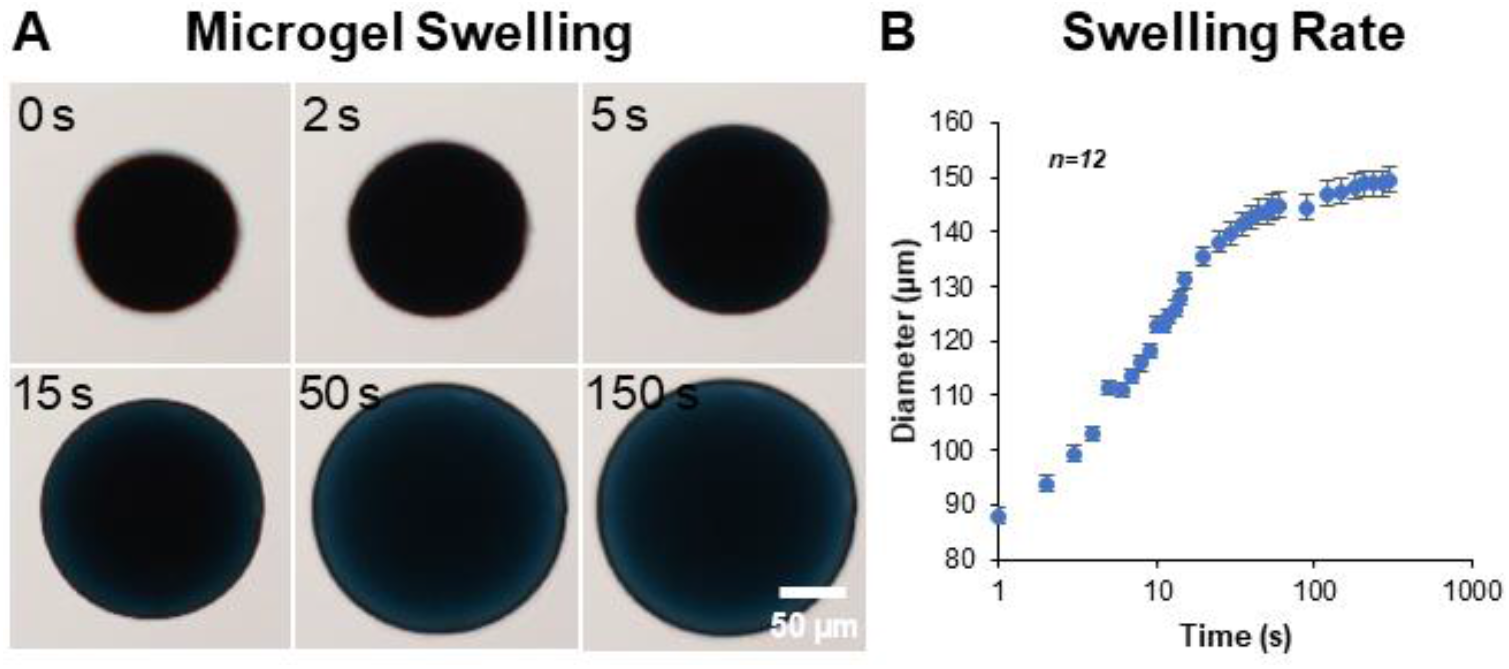
Microgel hydration and swelling. The microgels exhibited rapid swelling in an aqueous medium. A) Time-lapse bright-field images showing microgel being hydrated in PBS. B) The rate of hydration was rapid initially and plateaued within minutes to achieve an equilibrium state (n=12).

### 3.2. Licensing prolongs MSC immunomodulatory phenotype

We first investigated the concentration-dependent effects of two potent cytokines, IFNγ and TNFα, to sustain the immunomodulatory phenotype of adipose-derived MSC. A low (0.1 ng/mL) and high (100 ng/mL) concentration of the IFNγ and TNFα and a mixed treatment condition (10 ng/mL of each) were studied. MSC that express green fluorescence protein conjugated to the actin cytoskeleton (ACTb-EGFP) were used to monitor changes in morphology and viability longitudinally (**Fig. 3A**). No significant changes in morphology were seen in any treatment condition. However, the expression of *Nos2,* a key indicator of immunomodulatory phenotype, was significantly upregulated in all conditions (**Fig. 3B,** See **Table 1** for groupwise comparison). *Ptgs2,* a precursor to prostaglandin E2 and a potent immunomodulatory cytokine, was significantly upregulated in the 100 ng/mL TNFα treatment. Inflammatory cytokines *Ccl2* and *Il-6* were also significantly upregulated in most cases, with the most significant effect seen in groups with TNFα treatment. Interestingly, *Il-6* expression was not significantly affected by the IFNγ treatment *(p>0.05,* **Table 1**). Immunoregulatory and angiogenic proteins *Tgfβ* and *Lgals9were* significantly influenced by the 100 ng/mL TNFα treatment and combination groups, with *Lgals9* being downregulated and *Tgfβ* being upregulated. The TNFα treatment negatively influenced *Tnfα* expression, while no significant change was seen in the IFNγ treatment. The greatest increase in the most potent immunomodulatory factors *(Nos2* and *Ptgs2)* was seen in the TNFα treatment, but it also exhibited the highest upregulation of inflammatory cytokines (*Il-6* and *Ccl2).* IFNγ treatment, on the other hand, upregulated immunomodulatory factors while having modest effects on inflammatory factors. Hence, IFNγ is deemed more suitable for sustained licensing of adipose-derived MSC.

**Figure 3.**
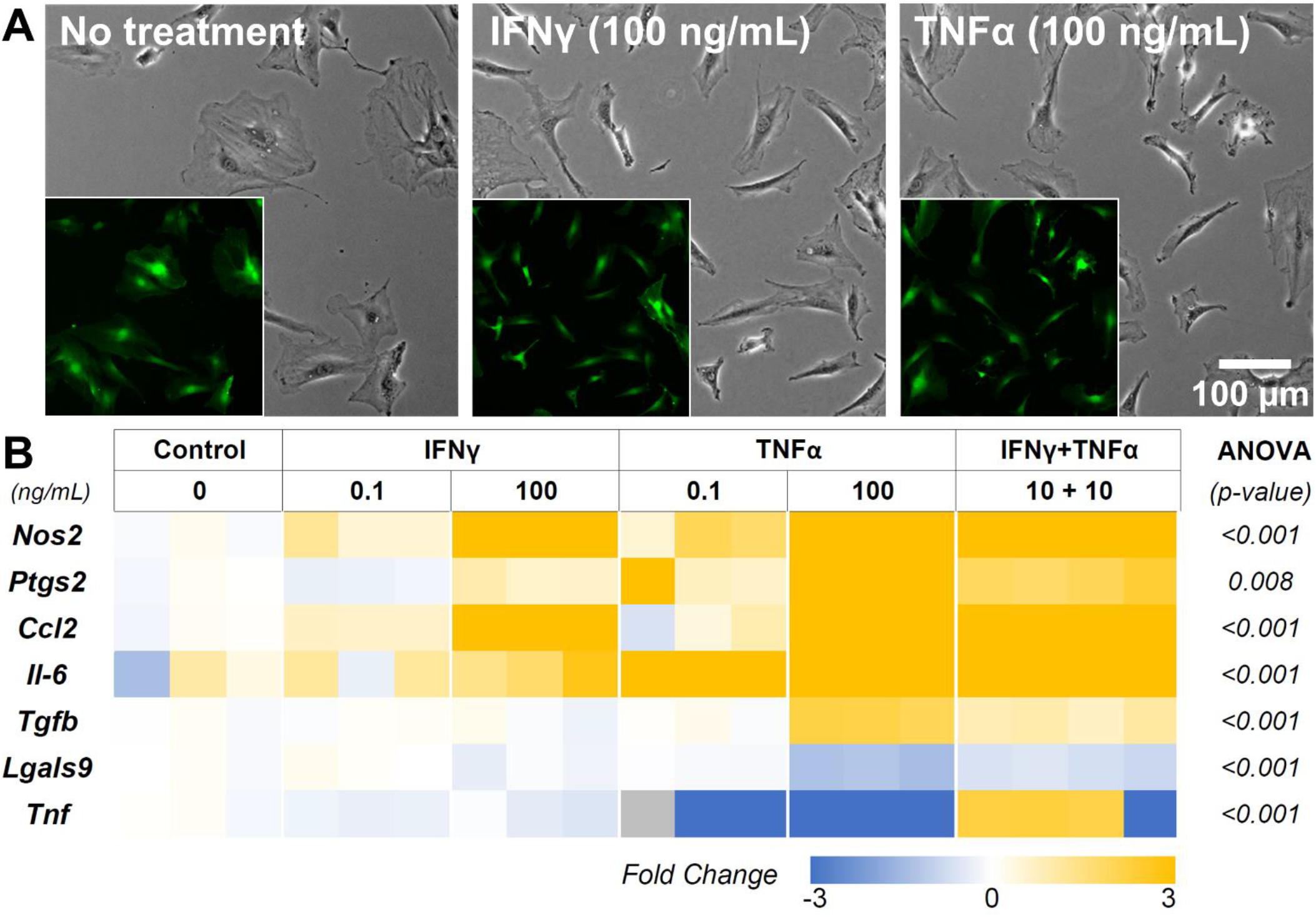
MSC Licensing and their Immunomodulatory Phenotype. **A)** Phase-contrast images of MSC control and treatment conditions after 12 hours. (Insets) Corresponding fluorescent images of the cells show minimal differences in morphology and viability. **B)** Heatmap of the MSC gene expression after 12 hours of licensing using low and high concentrations of IFNγ and TNFα. Heatmaps show fold change with respect to the housekeeping gene (Gapdh) and untreated control.

**Table 1.** MSC licensing through IFNγ and TNFα treatment. indicates p < 0.05

| Holm-Sidak Pairwise Comparison (vs. Control) (p-value) |  |  |  |  |  |  |
| --- | --- | --- | --- | --- | --- | --- |
| Genes | ANOVA (p-value) | IFNγ | | TNFα | | IFNγ+TNFα |
| (ng/mL) | - | 0.1 | 100 | 0.1 | 100 | 10 + 10 |
| *Nos2* | <0.001 | 0.024* | <0.001* | <0.001* | <0.001* | <0.001* |
| *Ptgs2* | 0.008 | 0.77 | 0.673 | 0.094 | 0.011* | 0.099 |
| *Ccl2* | <0.001 | 0.307 | <0.001* | 0.584 | <0.001* | <0.001* |
| *Il-6* | <0.001 | 0.605 | 0.274 | <0.001* | <0.001* | <0.001* |
| *Tgfb* | <0.001 | 0.923 | 0.751 | 0.95 | <0.001* | <0.001* |
| *Lgals9* | <0.001 | 0.546 | 0.99 | 0.357 | <0.001* | <0.001* |
| *Tnf* | <0.001 | 0.797 | 0.947 | <0.001* | 0.003* | 0.816 |

### 3.3. IFNγ licensing retains MSC phenotypes in a concentration-dependent manner

In-depth screening of IFNγ treatment with a range of concentrations (0.1, 1, 10, and 100 ng/mL) showed a concentration-depended increase in the expression levels of the key immunomodulatory markers (**Fig. 4A, Table 2**). The genes *Nos2, Ptgs2,* and *Ccl2* are upregulated in a concentration-dependent manner, with peak expression levels at 100 ng/mL treatment. Interestingly, *Il-6* expression was upregulated at 1 and 10 ng/mL, but no change was seen at 100 ng/mL. Likewise, *Tnfα* expression showed significant downregulation at 1 and 10 ng/mL, but no change was seen at 100 ng/mL. *Tnfa* and *Il-6* are potent proinflammatory cytokines that could negatively impact the regenerative milieu. Hence, we deemed concentrations from 10 and 100 ng/mL suitable for subsequent studies.

**Figure 4.**
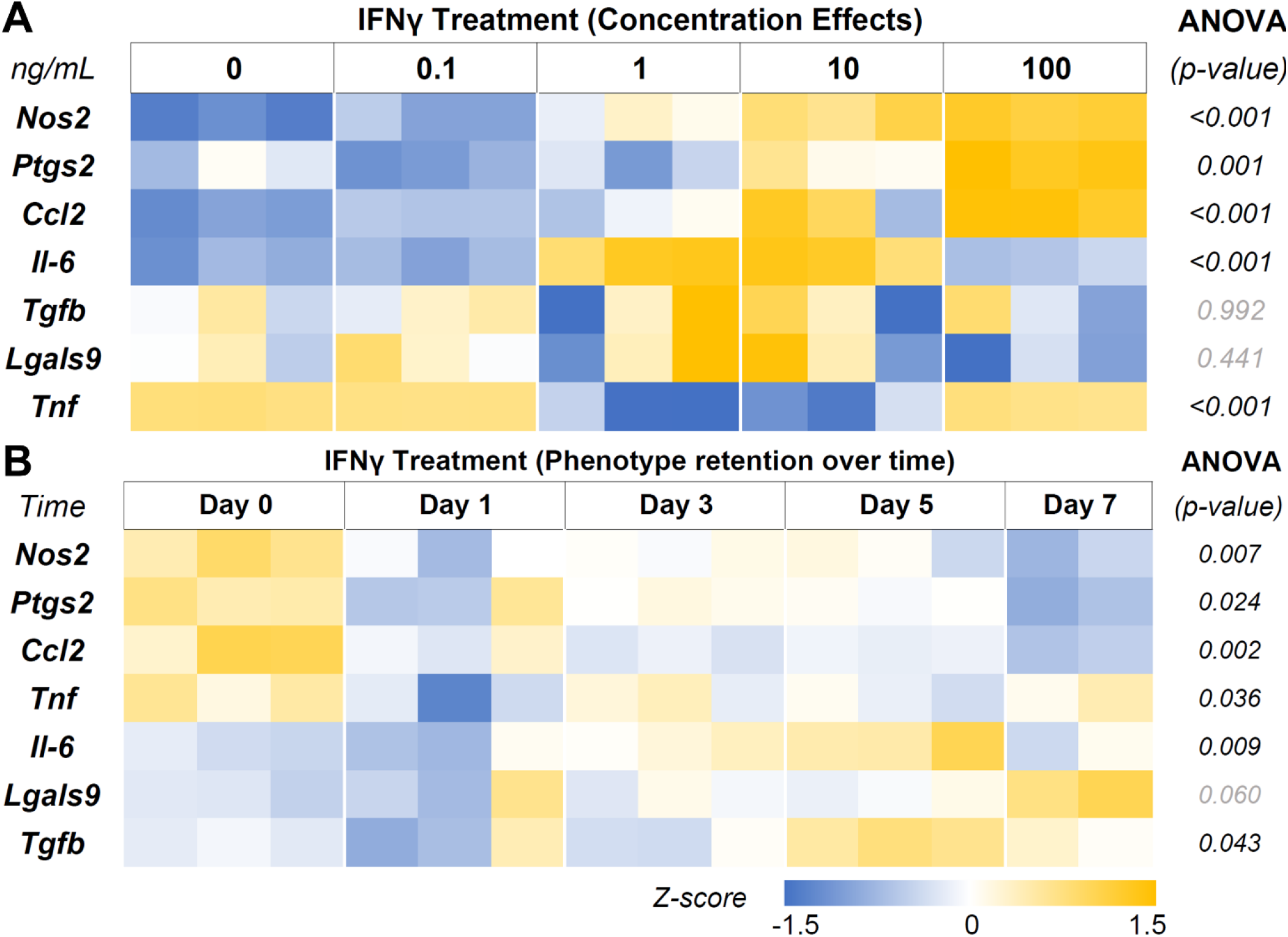
IFNγ Licensing and Retention of MSC Immunomodulatory Phenotype. A) Heatmap of the MSC gene expression after 24 hours of licensing using different concentrations of IFNγ shows a concentration-dependent effect. B) Heatmap of MSC licensed for 24 hours using 10 ng/mL IFNγ showing robust phenotype retention over time.

**Table 2.** IFNγ licensing is concentration-dependent – Groupwise comparison. N/A indicates no significance in ANOVA between groups. * indicate a significant change in gene expression.

| Holm-Sidak Pairwise Comparison (vs. Control) (*p*-value) |  |  |  |  |  |
| --- | --- | --- | --- | --- | --- |
| Genes | ANOVA<br>(*p*-value) | IFNγ | | | |
| (ng/mL) | - | 0.1 | 1 | 10 | 100 |
| **Nos2** | <0.001 | 0.014* | <0.001* | <0.001* | <0.001* |
| **Ptgs2** | 0.001 | 0.129 | 0.367 | 0.164 | <0.001* |
| **Ccl2** | <0.001 | 0.283 | 0.136 | 0.011* | <0.001* |
| **Il-6** | <0.001 | 0.53 | <0.001* | <0.001* | 0.168 |
| **Tgfb** | 0.992 | N/A | N/A | N/A | N/A |
| **Lgals9** | 0.441 | N/A | N/A | N/A | N/A |
| **Tnf** | <0.001 | 0.872 | <0.001* | <0.001* | 0.979 |

Then we investigated the retention of the licensed MSC phenotype over 7 days after the initial treatment. The MSC were treated with 10 ng/mL of IFNγ for 24 hours, washed thoroughly to remove all cytokines, and monitored for 7 days (**Fig. 4B, Table 3**). *Nos2,* one of the critical factors of the licensed phenotype, was downregulated after 24 hours. In contrast, *Ptgs2* expression remained consistent until day 7, when it was significantly downregulated, indicating a sustained MSC phenotype. *Ccl2* expression dropped within 24 hours and continued to drop until day 7. *Tnfα* expression, on the other hand, showed downregulation after 24 hours but bounced back to licensed levels. *Il-6* expression remained unchanged until day 7, while *Lgals9 and Tgfb* did not show significant changes. Overall, the licensed phenotype of the MSC were partially retained for a week after licensing signals were removed.

**Table 3.** MSC phenotype retention after IFNγ licensing pairwise comparison. N/A indicates no significance in ANOVA between groups. * indicate a significant change in gene expression.

| Holm-Sidak Pairwise Comparison (vs. Control) (*p*-value) |  |  |  |  |  |
| --- | --- | --- | --- | --- | --- |
| Genes | ANOVA<br>(*p*-value) | Day 1 | Day 3 | Day 5 | Day 7 |
| **Nos2** | 0.007 | 0.009* | 0.023* | 0.023* | 0.004* |
| **Ptgs2** | 0.024 | 0.077 | 0.151 | 0.132 | 0.009* |
| **Ccl2** | 0.002 | 0.003* | 0.002* | 0.003* | <0.001* |
| **Tnfa** | 0.036 | 0.019* | 0.546 | 0.199 | 0.614 |
| **Il-6** | 0.009 | 0.678 | 0.150 | 0.776 | 0.011* |
| **Lgals9** | 0.060 | N/A | N/A | N/A | N/A |
| **Tgfb** | 0.043 | 0.774 | 0.871 | 0.078 | 0.679 |

### 3.4. MSC seeded on licensing microgels exhibit a sustained immunomodulatory phenotype

MSC seeded on the microgels rapidly attach and spread on the surface and fully cover them within 24 hours (**Fig. 5A**). MSC seeded on IFNγ-loaded microgels showed a similar morphology and viability and sustained a licensed phenotype for 7 days with a consistent expression of *Nos2* over time (**Fig. 5B, Table 4**). The expression pattern for both the licensing microgels and the bolus treatment (IFNγ supplied in the media) groups was similar over 7 days, with minor differences in *Ccl2* and *Nos2* expression on days 1 and 7, respectively. Interestingly, both groups exhibited downregulation in *Ptgs2, Lgals9,* and *Tgfβ* on day 3 relative to the no treatment control, indicating the phenotype similarities between the treatment groups. *Il-6* was significantly downregulated on days 1 and 3 in the microgel group and only on day 3 in the bolus treatment group. No significant change in *Tnfα* expression was noticed in any conditions. Overall, the microgels enhanced and protracted the immunomodulatory phenotype of MSC as effectively as the bolus group without needing constant replenishment of the licensing factors.

**Figure 5.**
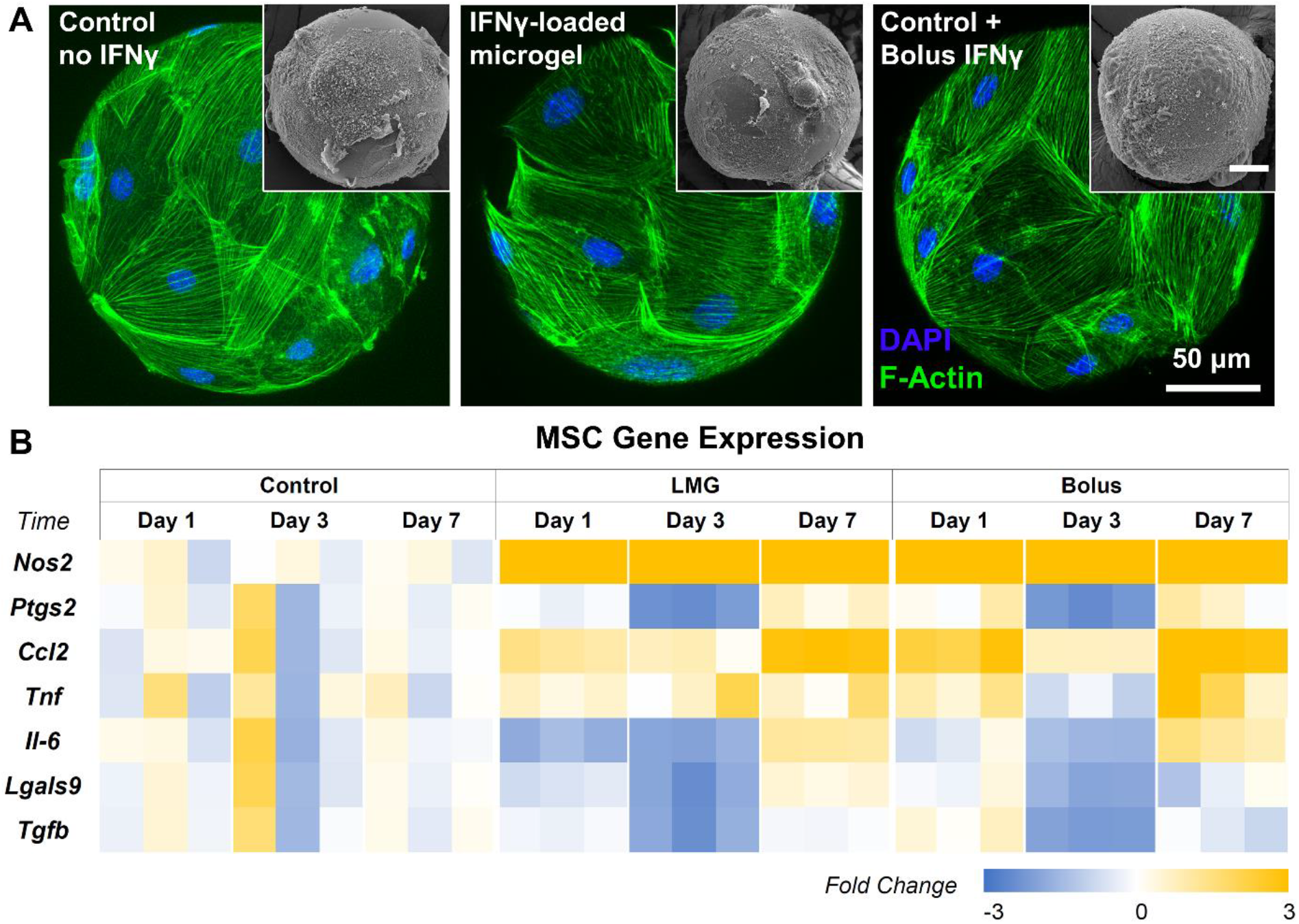
Gene Expression of MSC Seeded on licensing Microgels. **A)** Confocal Z-stacks of microgels seeded with MSC (Control microgels, IFNγ-loaded microgels, and IFNγ bolus treatment). (Insets) Corresponding SEM images of microgels. **B)** Heatmap of MSC gene expression of inflammatory and anti-inflammatory MSC phenotypes. Heatmaps show fold change with respect to the housekeeping gene (Gapdh) and no treatment control group.

**Table 4.** Pairwise comparison of all Time Points within Conditions and all Conditions within Time Points. N/A indicates no significance in ANOVA between groups. * indicate a significant change in gene expression.

| Holm-Sidak Pairwise Comparison (Time Point within Condition) (*p*-value) |  |  |  |  |  |  |  |  |  |
| --- | --- | --- | --- | --- | --- | --- | --- | --- | --- |
|  | Control |  |  | MGL |  |  | Bolus |  |  |
| Genes | Day 1 vs Day 3 | Day 1 vs Day 7 | Day 3 vs Day 7 | Day 1 vs Day 3 | Day 1 vs Day 7 | Day 3 vs Day 7 | Day 1 vs Day 3 | Day 1 vs Day 7 | Day 3 vs Day 7 |
| *Nos2* | >0.999 | >0.999 | >0.999 | 0.036* | 0.035* | 0.862 | 0.004* | 0.095 | 0.12 |
| *Ptgs2* | >0.999 | >0.999 | >0.999 | 0.002* | 0.198 | <0.001* | <0.001* | 0.777 | <0.001* |
| *Ccl2* | >0.999 | >0.999 | >0.999 | 0.274 | 0.014* | 0.002* | 0.014* | 0.204 | 0.001* |
| *Tnf* | NA | NA | NA | NA | NA | NA | NA | NA | NA |
| *Il-6* | >0.999 | >0.999 | >0.999 | 0.737 | <0.001* | <0.001* | 0.061 | 0.042* | <0.001* |
| *Lgals9* | >0.999 | >0.999 | >0.999 | 0.043* | 0.086 | 0.001* | 0.016* | 0.4 | 0.067 |
| *Tgfb* | >0.999 | >0.999 | >0.999 | 0.004* | 0.886 | 0.004* | <0.001* | 0.083 | 0.008* |

| Holm-Sidak Pairwise Comparison (Condition within Time Point) (*p*-value) |  |  |  |  |  |  |  |  |  |
| --- | --- | --- | --- | --- | --- | --- | --- | --- | --- |
|  | Day 1 |  |  | Day 3 |  |  | Day 7 |  |  |
| Genes | Control vs MGL | Control vs Bolus | MGL vs Bolus | Control vs MGL | Control vs Bolus | MGL vs Bolus | Control vs MGL | Control vs Bolus | MGL vs Bolus |
| *Nos2* | <0.001* | <0.001* | 0.117 | <0.001* | <0.001* | 0.54 | <0.001* | <0.001* | 0.012* |
| *Ptgs2* | 0.785 | 0.73 | 0.702 | 0.001* | 0.002* | 0.997 | 0.661 | 0.547 | 0.958 |
| *Ccl2* | 0.083 | 0.001* | 0.045* | 0.517 | 0.498 | 0.799 | <0.001* | <0.001* | 0.669 |
| *Tnf* | NA | NA | NA | NA | NA | NA | NA | NA | NA |
| *Il-6* | 0.037* | 0.598 | 0.074 | 0.018* | 0.041* | 0.568 | 0.144 | 0.174 | 0.921 |
| *Lgals9* | 0.518 | 0.977 | 0.647 | 0.007* | 0.012* | 0.665 | 0.454 | 0.658 | 0.336 |
| *Tgfb* | 0.676 | 0.626 | 0.501 | 0.002* | 0.001* | 0.848 | 0.784 | 0.731 | 0.76 |

### 3.5. Licensed MSC modulate phenotype of inflammatory macrophages

To validate the immunomodulatory effects, MSC seeded in 2D tissue culture plastics and microgels were licensed with bolus IFNγ and cocultured with proinflammatory M1-like macrophages in trans-well plates, as shown (**Fig. 6A**). All cells were washed thoroughly in PBS to remove all biochemical factors before initiating the coculture. Serum-free media was used for coculture studies. After 24 hours of coculture, MSC and macrophage samples were collected and analyzed individually. M1 macrophage monocultures served as controls. In all coculture conditions, the MSC modulated macrophages from an M1-like phenotype toward a proreparative M2-like phenotype (**Fig. 6B, Table. 5**). In general, the MSC coculture significantly upregulated M2 markers *Mrc1, Pparγ,* and *Arg1* and downregulated M1 markers *Il-6, Cxcl2, Tnfα,* and *Il-1β* in macrophages. But, *Pparγ* expression was upregulated at a relatively higher level when MSC were licensed with 1 ng/mL than at 100 ng/mL IFNγ concentration. Likewise, *Cxcl2* was downregulated more prominently by MSC licensed at 100 ng/mL IFNγ than the other treatment conditions (For pairwise comparisons, see **Table. 5**).

**Figure 6.**
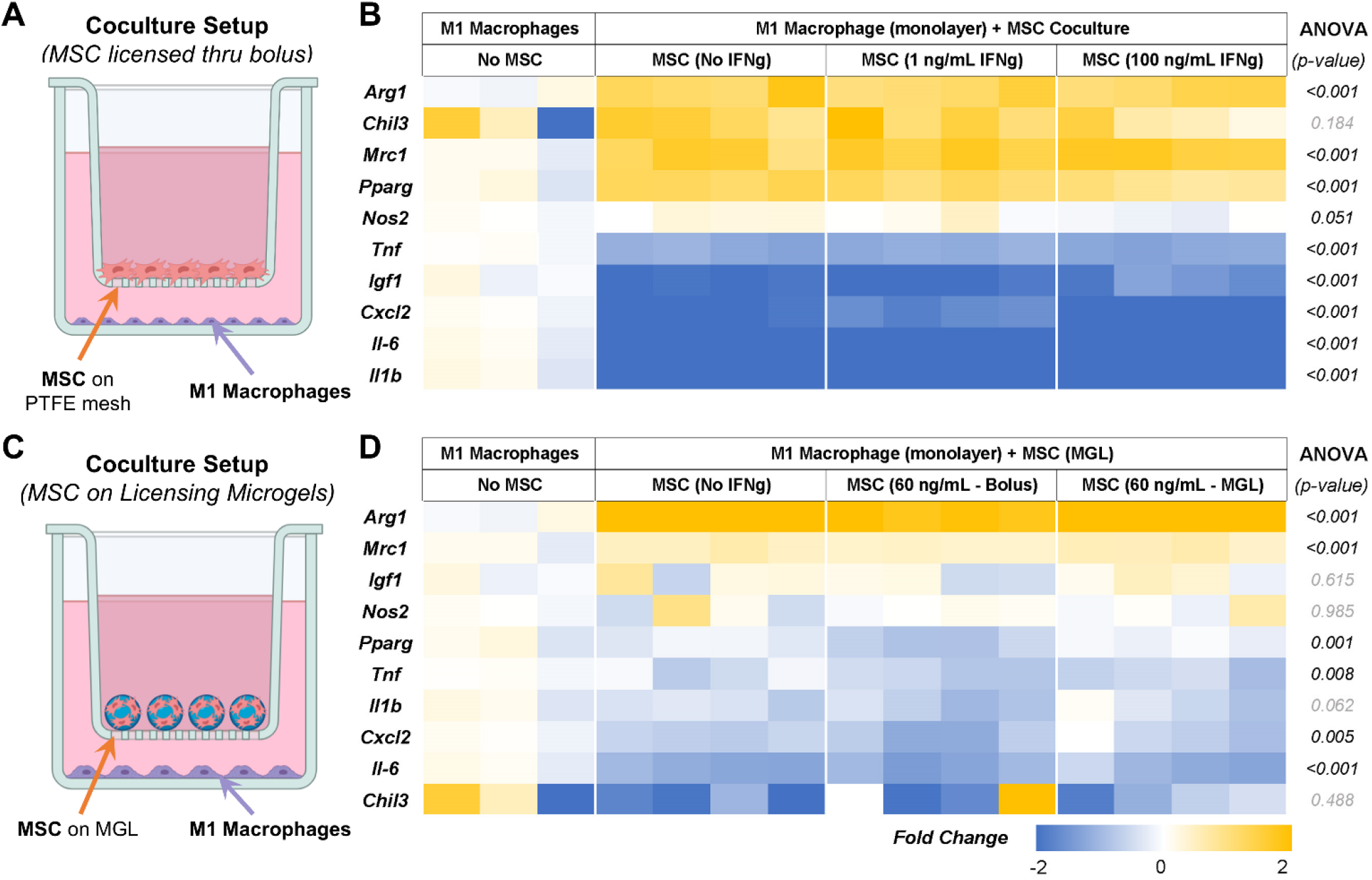
MSC and M1 Macrophage Cocultures. **A)** Schematic of the coculture setup to evaluate the MSC paracrine effects on M1-like macrophages. **B)** Heatmap showing the expression levels of proinflammatory and prohealing genes by macrophages after 24 hours of coculture with licensed MSC. **C)** Schematic of the coculture setup showing MSC seeded on microgels in the trans-well and M1 macrophages on the bottom well. **D)** Heatmap showing the expressional levels of proinflammatory and prohealing genes by macrophages after 24 hours of coculture with MSC on microgels. All fold changes are with respect to the housekeeping gene, Gapdh, and quiescent control MSC.

**Table 5.** Pairwise comparison between all treatment conditions. N/A indicates no significance in ANOVA between groups. * indicate a significant change in gene expression.

| Holm-Sidak Pairwise Comparison (Between Groups) (*p-value*) |  |  |  |  |  |  |  |
| --- | --- | --- | --- | --- | --- | --- | --- |
| Genes | ANOVA<br>(*p-value*) | M1 vs<br>Control | M1 vs 1 ng<br>IFNγ | M1 vs 100 ng<br>IFNγ | Control<br>vs 1 ng<br>IFNγ | Control<br>vs 100 ng<br>IFNγ | 1 ng vs<br>100 ng<br>IFNγ |
| *Arg1* | <0.001 | <0.001* | <0.001* | <0.001* | 0.809 | 0.846 | 0.770 |
| *Chil3* | 0.184 | N/A | N/A | N/A | N/A | N/A | N/A |
| *Mrc1* | <0.001 | <0.001* | <0.001* | <0.001* | 0.469 | 0.480 | 0.832 |
| *Ppary* | <0.001 | <0.001* | <0.001* | <0.001* | 0.439 | 0.026* | 0.071 |
| *Nos2* | 0.051 | N/A | N/A | N/A | N/A | N/A | N/A |
| *Tnfa* | <0.001 | <0.001* | <0.001* | <0.001* | 0.745 | 0.553 | 0.496 |
| *Igf1* | <0.001 | <0.001* | <0.001* | <0.001* | 0.643 | 0.027* | 0.018* |
| *Cxcl2* | <0.001 | <0.001* | <0.001* | <0.001* | 0.004* | 0.110 | <0.001* |
| *Il-6* | <0.001 | <0.001* | <0.001* | <0.001* | 0.812 | 0.081 | 0.084 |
| *Il1β* | <0.001 | <0.001* | <0.001* | <0.001* | 0.286 | 0.171 | 0.584 |

Then we investigated the effects of MSC licensed using IFNγ sequestered on microgels on M1 macrophages and how they compare to other treatment groups. The microgels seeded with MSC were separated from the M1 macrophages using transwells, as shown (**Fig. 6C**). We used 1/10^th^ of the maximum loading capacity of the microgels (60 ng of IFNγ /mg of lyophilized microgels) and evaluated their potential in modulating MSC phenotype and function using coculture studies. We compared the effects of the licensing microgels to an equivalent bolus media (60 ng/mL of IFNγ) treatment that is renewed every other day. Untreated MSC served as controls. In all coculture conditions, the MSC were seeded on the microgels, as shown (**Fig. 6C**). As seen with monolayer cultures, the MSC seeded on all microgels groups modulated the M1 macrophages towards a proreparative M2-like phenotype. Notably, the IFNγ sequestered microgels outperformed both the untreated control and bolus treatment in downregulating the inflammatory cytokine *Tnfα* in macrophages (**Fig. 6D**). Similar levels of modulatory effects were seen in terms of upregulating *Arg1* and *Mrc1* and downregulating *Cxcl2* and *Il-6* between the bolus and the IFNγ sequestered microgel groups. Overall, these data show the notable effects of the IFNγ sequestered microgels in enabling and sustaining the MSC immunomodulatory phenotype in macrophage cocultures.

**Table 6.**
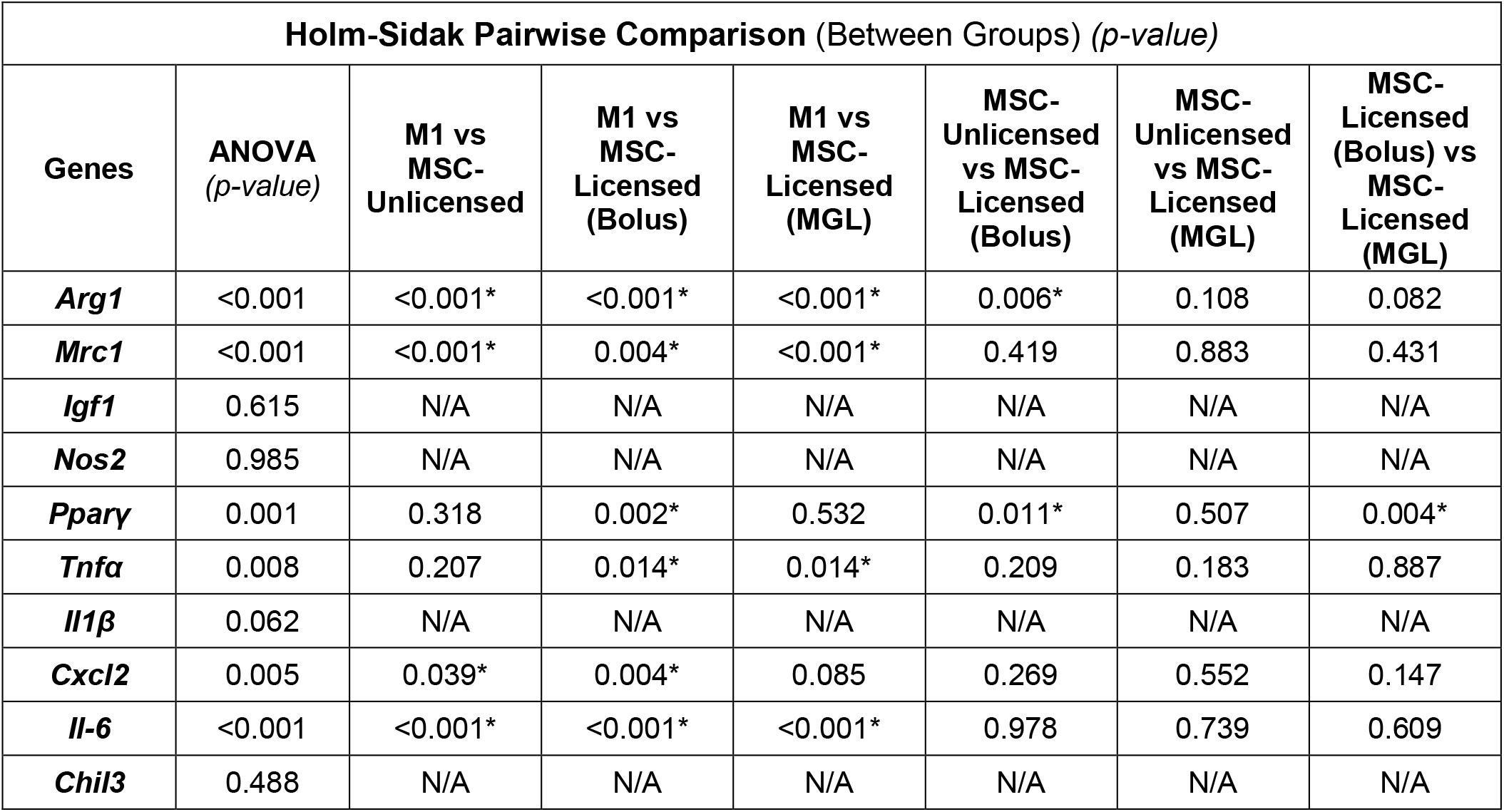
Pairwise comparison between all treatment conditions. N/A indicates no significance in NOVA between groups. * indicate a significant change in gene expression.

### 3.6. Microgels support MSC osteogenesis in the absence of IFNγ

To evaluate the long-term outcomes of the microgel cultures, we seeded the MSC in microgels and maintained them in growth (CNT) and osteogenic (OST) conditions for 21 days and evaluated their gene expression. In both the culture conditions, *Ibsp, Runx2, Col1a1,* and *Sp7* were upregulated (**Fig. 7A, Table 7**). Interestingly, *Alp,* an early-stage osteogenesis marker, was expressed at a higher level in the growth conditions than in osteogenic conditions. The enzymatic assay also revealed relatively higher ALP activity in microgels cultured in the growth conditions MSC compared to osteogenic conditions (**Fig. 7B**). Further, *Spp1* (aka osteopontin), a later-stage osteogenic marker, was expressed at lower levels in the growth media compared to the osteogenic media. Calcium deposition was noticeable in media conditions, with higher rates in the osteogenic conditions in the later stages (days 14 and 21). Overall, the microgels provide a suitable surface for osteogenic differentiation of MSC in the absence of IFNγ.

**Figure 7.**
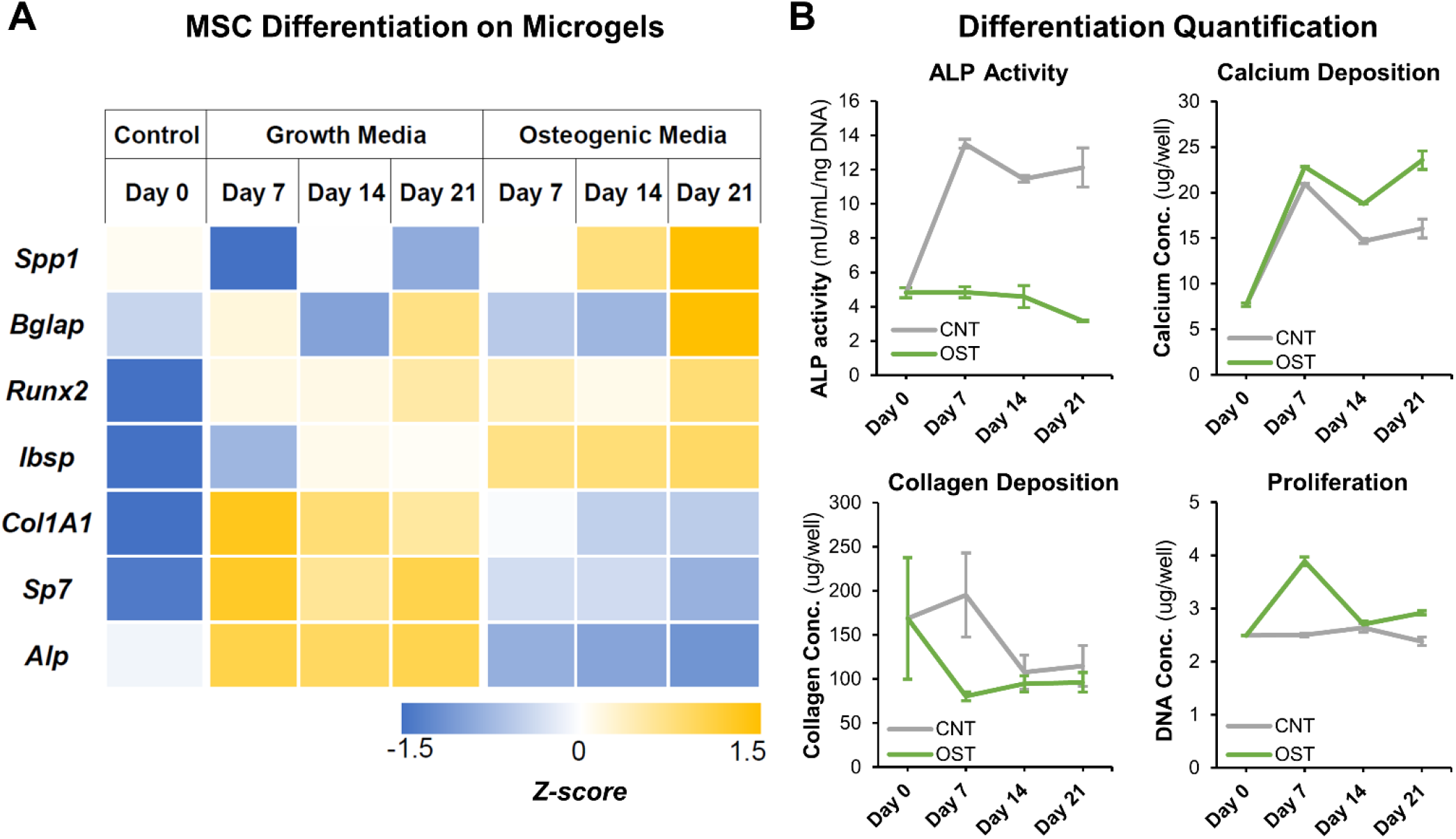
Osteogenic Gene Expression and Characterization. **A)** Fold change of common osteogenic markers when MSC are seeded on microgels with growth and osteogenic differentiation media. **B)** Quantification of ALP activity and calcium deposition.

**Table 7.**
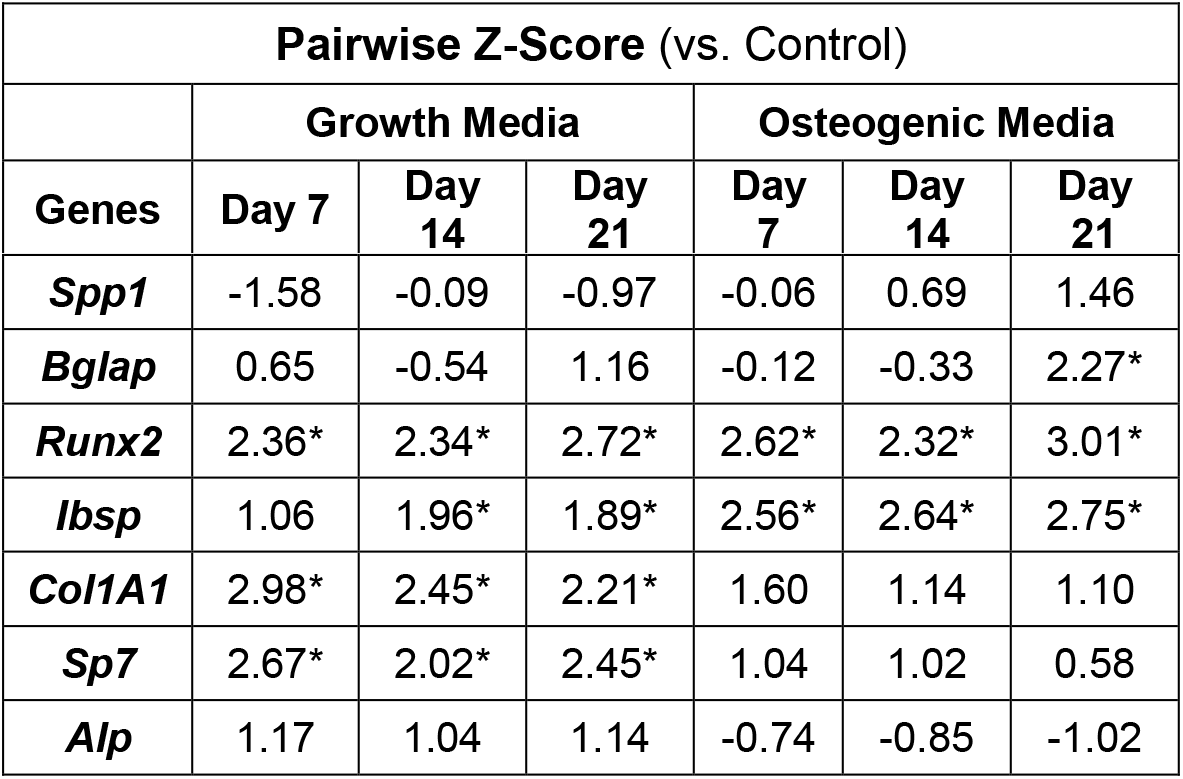
Pairwise comparison of all treatment conditions with control. p(−1.65>Z>1.65) <0.05

## 4. Discussion

MSC can be derived from various adult tissue sources and exhibit self-renewal and multipotent capacities. They are the sensors of the body’s immune response and possess immune modulatory functions. They can promote inflammation when the immune response is underactive and restrain the inflammatory response when it is overactive. The immunomodulatory effects of MSC vary depending on their tissue source and activation state. The major immunomodulatory factors secreted by MSC include NO, PGE2, TGFβ, Galactin 9, and TSG6^8^. In our studies, we freshly isolated MSC from inguinal fat pads (adipose-derived MSC) because of the clinical relevance of the tissue source. When the MSC were licensed with IFNγ or TNFα, they consistently upregulated a set of markers, including *Nos2, Ccl2,* and *Il-6*. *Nos2* was the most highly upregulated gene which results in the production of nitric oxide and subsequent suppression of T-cell proliferation^22^. *Ccl2,* aka monocyte chemoattractant protein-1, is generally responsible for mobilizing monocytes, memory T lymphocytes, and natural killer cells to initiate inflammatory response^23^. But, studies using murine bone marrow MSC found that *Ccl2* polarizes inflammatory macrophages toward regenerative (IL10-secreting) macrophages and treats inflammatory colitis^24^. Notably, when TNFα alone was used for licensing, *Il-6* was highly upregulated in MSC. IL-6 is an inflammatory cytokine released in response to infection or tissue damage, triggering an inflammatory response from regulatory cells. They also play a crucial role in directing Th17 differentiation on the Th17/Treg axis^25^. Further, the expression of *Ptgs2,* necessary for PGE2 enzyme synthesis, is positively influenced only when MSC were treated with IFNγ and not with TNFα treatment. Due to the elevated expression of *Il-6* by MSC in all treatment formulations containing TNFα, IFNγ was chosen as the best licensing candidate for our studies. IFNγ showed concentration-dependent licensing effects on MSC, with low concentrations (<10 ng/mL) showing minimal licensing effects with high expression of *Il-6.* But at medium to high concentrations (10-100 ng/mL), they promoted a robust immunomodulatory MSC phenotype where *Ptgs2* is upregulated, and *Tnfα* is downregulated. Over time, some licensed phenotypes, notably, *Nos2* and *Ccl2* expressions, diminish if the media is not replenished with IFNγ. Then we tested the ability of the licensing microgels to prolong a licensed-MSC phenotype.

The unique physical and chemical properties of gelatin microgels make them versatile carriers for cell and growth factor delivery. They can form complexes with different drugs, cytokines, and growth factors and therefore widely used in drug delivery and tissue engineering applications^26^. The tunable properties of the gelatin matrix, such as the crosslinking density and the isoelectric point, enable the optimization of degradation and drug delivery kinetics. The microscale nature of the microgels we fabricated allows them to be injected minimally invasively using hypodermic needles and conformally fill various tissue defects. Our studies show that the enzymatic degradation of microgels is concentration-dependent and hence can allow delivery of sequestered factors in a bioresponsive manner, particularly in inflammatory conditions like osteoarthritis, where there are elevated levels of matrix metalloproteases (MMPs) during flare-ups^27^. Hence the microgels can titer a therapeutic response based on the disease state. Further, we show that the degradation of the microgels and the release of sequestered IFNγ from the microgels are linearly correlated, indicating a homogenous polymer density and cytokine sequestration. For our cell culture studies, the average size of the microgels was kept at 200 μm in diameter, large enough to allow MSC to attach and spread on the surface and injectable when needed. The high hydration and the swelling rate (>400% volume increase) of the microgels also show cytocompatibility, which can be tailored by varying the crosslinking density. The MSC readily attached and spread on the surface of the microgels. As expected, the loaded microgels were able to sustain the established licensed MSC characteristics over 7 days at a similar level to that of the bolus control. The IFNγ sequester by the microgels is slowly released as the MSC, known to secrete MMP^28^, degrades the gelatin. This sustained licensed state of MSC is critical to addressing chronic or reoccurring inflammation and can help improve the efficacy of MSC therapies.

To confirm the therapeutic effects of the licensed MSC, coculture studies were performed using IFNγ licensed MSC and M1-like macrophages. As expected, the licensed MSC were able to modulate the inflammatory macrophage phenotype toward a reparative M2-like phenotype. Especially when MSC were licensed using microgels, they upregulated *Arg1, Mrc1,* and *Igf1* in macrophages more robustly compared to monolayer cultures. These enhancements can be attributed to the tissue-like microenvironment provided by the microgels. It is known that substrate mechanics influence MSC phenotype, including their immunomodulatory capabillites^29^. In addition, the licensed MSC significantly downregulated *Tnfα* compared to the unlicensed MSC. In the absence of any licensing cytokines, the microgels also supported MSC osteogenesis, presumably due to their spherical morphology, which is known to promote MSC osteogenesis^30, 31^.

In conclusion, we show that microgels efficiently sequester IFNγ and release it upon degradation while readily supporting MSC attachment. We also show that the localized release of IFNγ licenses the MSC efficiently and further confirm their immunomodulatory phenotype through macrophage coculture studies. Overall, licensing MSC using the microgel system is a simple and effective tool to sustain the immunomodulatory properties of MSC and enhance their therapeutic applications in various instances.

## 5. Acknowledgments

Research reported in this publication was supported in part by the National Institute of Arthritis and Musculoskeletal and Skin Diseases (NIAMS) Award Number R21AR078447, National Institute of General Medical Sciences (NIGMS) of the National Institutes of Health under Award Numbers P20GM130456 and P20GM103436-20 (KY IDeA Networks of Biomedical Research Excellence), National Center for Research Resources and the National Center for Advancing Translational Sciences of the National Institutes of Health under Award Number UL1TR001998, and Orthopedic Trauma Association (OTA, Grant Number: 6889). The content is solely the responsibility of the authors and does not necessarily represent the official views of the National Institutes of Health or other grant funding agencies.

